# Lexical selection with competing distractors: Evidence from left temporal lobe lesions

**DOI:** 10.1101/085670

**Authors:** Vitória Piai, Robert T. Knight

**Author notes:** Author note: Correspondence concerning this article should be addressed to Vitória Piai, Donders Centre for Cognition, Radboud University, Nijmegen, the Netherlands. The authors are grateful to the patients and their families, as well as to the other volunteering participants for taking part in this study. We would like to thank Donatella Scabini for patient delineation, Brian Curran and Callum Dewar for lesion reconstruction, the members of the Center for Aphasia and Related Disorders at the VANCHCS in Martinez, CA, for neuropsychological testing, and Ardi Roelofs, Peter Indefrey, and Kristoffer Dahlslätt for helpful discussions. This work is supported by grants from the Netherlands Organization for Scientific Research (446-13-009 to V.P.) and National Institutes of Health (NINDS R37 NS21135 to R.T.K).

## Abstract

According to the competition account of lexical selection in word production, conceptually driven word retrieval involves the activation of a set of candidate words in left temporal cortex, and competitive selection of the intended word from this set, regulated by frontal cortical mechanisms. However, the relative contribution of these brain regions to competitive lexical selection is uncertain. In the present study, five patients with left prefrontal cortex lesions (overlapping in ventral and dorsal lateral cortex), eight patients with left lateral temporal cortex lesions (overlapping in middle temporal gyrus), and 13 matched controls performed a picture-word interference task. Distractor words were semantically related or unrelated to the picture, or the name of the picture (congruent condition). Semantic interference (related vs unrelated), tapping into competitive lexical selection, was examined. An overall semantic interference effect was observed for the control and left-temporal groups separately. The left-frontal patients did not show a reliable semantic interference effect as a group. The left-temporal patients had increased semantic interference in the error rates relative to controls. Error distribution analyses indicated that these patients had more hesitant responses for the related than for the unrelated condition. We propose that left middle temporal lesions affect the lexical activation component, making lexical selection more susceptible to errors.

## Introduction

Selecting words for speaking is a competitive process (Levelt, Roelofs, & Meyer, 1999; Roelofs, 1992; Spalek, Damian, & Bölte, 2013), involving not only core language processes, such as lexical retrieval, but also mechanisms for attentional control (Roelofs & Piai, 2011). However, the relative contribution of different brain regions to competitive lexical selection in word production is uncertain. According to the competition view, conceptually driven word retrieval involves the activation of a set of candidate words in left middle temporal cortex. Competitive selection of the intended word from this set is regulated by frontal cortical mechanisms (Roelofs & Piai, 2011).

In picture-word interference, participants name pictures presented along with a distractor word, with performance depending on the relationship between the picture name and the distractor word. If the distractor is incongruent with and unrelated to the picture name (e.g., picture: pig, distractor “chair”), picture naming is more difficult relative to congruent distractors (e.g., picture: pig, distractor “pig”, Piai, Roelofs, Acheson, & Takashima, 2013). If the distractor is from the same semantic category as the picture (e.g., picture: pig, distractor “cow”), picture naming is more difficult relative to unrelated distractors (Glaser & Düngelhoff, 1984). Figure 1 gives an example of each picture-word distractor condition. The semantic relationship between the distractor and the picture makes the distractor a stronger competitor for the picture name relative to a semantically unrelated word (Roelofs, 1992, 2003). Thus, the semantic interference effect has been a key focus for investigating the competitive nature of lexical selection in word production.

**Figure 1.**
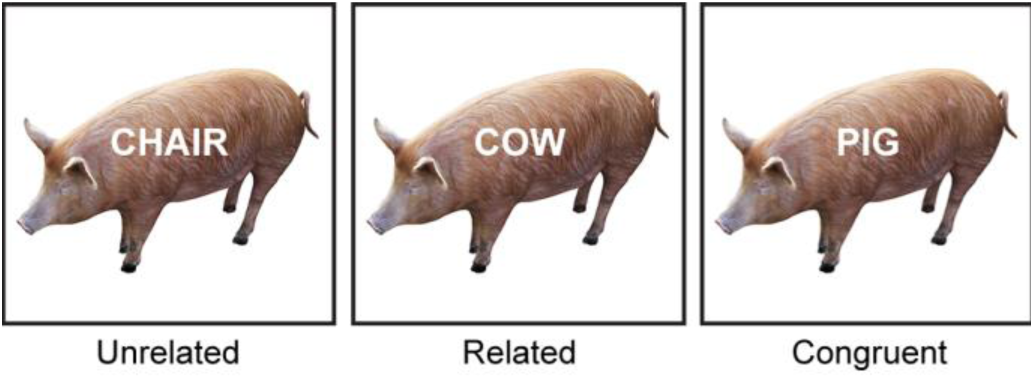
An example of each distractor condition. Materials were obtained from the BOSS database (Brodeur et al., 2010). Pictures are shown in black and white in the figure, but were shown in color during the experiment.

Previous neuroimaging studies have provided converging evidence for the involvement of two brain areas in the semantic interference effect from distractors: left temporal and frontal cortex. Activity in the left-temporal cortex has been shown to decrease with semantically related relative to unrelated distractors (de Zubicaray, Hansen, &McMahon, 2013; Piai et al., 2013; Piai, Roelofs, Jensen, Schoffelen, & Bonnefond, 2014). This decreased activity has been interpreted in terms of semantic priming between picture and distractor, thus reflecting the lexical activation mechanism (Piai et al., 2013, 2014). By contrast, activity in frontal cortex, in particular superior frontal gyrus and anterior cingulate cortex, increases for semantically related relative to unrelated distractors (Piai et al., 2013, 2014; see also de Zubicaray, Wilson, McMahon, & Muthiah, 2001). This increased frontal activity has been interpreted as reflecting the top-down control signal over lexical representations in the temporal cortex.

The critical involvement of the lateral prefrontal cortex (PFC) to the resolution of competition in word production has been found in some picture-naming tasks with lesion-symptom investigations (Riès, Greenhouse, Dronkers, Haaland, & Knight, 2014; Schnur et al., 2009). However, to date the contribution of this brain area to the semantic interference effect from distractor words has remained unclear (de Zubicaray et al., 2013; Piai, Riès, & Swick, 2016; Piai et al., 2013, 2014). Piai et al. (2016) examined picture-word interference in six patients with 100% lesion overlap in the ventrolateral PFC. On the group level, no consistent semantic interference effect was found. Descriptively, three patients showed semantic interference and three patients showed semantic facilitation.

In the present study, patients with left-lateral temporal or frontal lesions named pictures while ignoring semantically related, unrelated, and congruent visual distractors. We maximized the amount of competition exerted by the distractors. Firstly, distractor and picture were presented simultaneously. Secondly, congruent distractors were included. In the color-word Stroop task, the presence of congruent trials (e.g., “red” displayed in red ink) adds relevance to the task dimension (here, word reading) that should otherwise be ignored (Lowe & Mitterer, 1982). This manipulation induces a larger Stroop interference effect. We reasoned that a similar attentional mechanism could be at play in picture-word interference. Finally, distractor words also appeared as pictures in the experiment (i.e., they were part of the response set). For example, “pig” appeared as a picture in some trials but as a distractor in other trials. The increased interference from response-set members has been shown for tasks such as color-word Stroop (Klein, 1964; Lamers, Roelofs, & Rabeling-Keus, 2010) and picture-word interference (Piai, Roelofs, & Schriefers, 2012). In Piai et al. (2016), the distractor words were not in the response set nor was the congruent condition included. Thus, the materials of Piai et al. (2016) may have been too weak to induce semantic interference, explaining why their patients with left-ventrolateral PFC lesions did not show an abnormally large semantic interference effect. Regarding the left temporal cortex, to the best of our knowledge, no picture-word interference study has been published with patients with well-characterized left-temporal lesions. So the critical role of the left-temporal cortex to the semantic interference effect of distractor words is largely unknown.

## Method

The study protocol was approved by the University of California, Berkeley Committee for Protection of Human Subjects, following the declaration of Helsinki. Participants gave written informed consent and received monetary compensation for participating.

### Participants

Thirteen patients participated. Eight had a main lesion in the left lateral-temporal cortex (one female; median age = 70, mean = 67, sd = 8, range = 50-74; mean years of education = 17) and five had a main lesion in the left PFC (one male; median/mean age = 64 sd = 9, range = 53-73; mean years of education = 16). Patients were pre-morbidly right handed. Information on the patients’ lesions and language ability are shown in Tables 1 and 2. Additionally, 13 right-handed controls participated, each matched closely for gender, age, and years of education to their matched patient within ±4 years of age and ±2 years of education (five females; median age = 68, mean = 65, sd = 7.6, range = 50-74, *t*(12) = 0, *p* = 1; mean years of education = 16.6, *t*(12) < 1, *p* = .695). All participants were native speakers of American English. None of the participants had a history of psychiatric disturbances, substance abuse, multiple neurological events, or dementia.

**Table 1.** Individual lesion volume and percent damage to the left inferior frontal gyrus (IFG), middle frontal gyrus (MFG), superior frontal gyrus (SFG), superior temporal gyrus (STG), and middle temporal gyrus (MTG).

| Patient | Lesion volume | IFG | MFG | SFG | STG | MTG |
| --- | --- | --- | --- | --- | --- | --- |
| Left temporal lobe lesions |  |  |  |  |  |  |
| 1 | 18.32 | 0 | 0 | 0 | 34 | 23.6 |
| 2 | 93.75 | 0 | 0 | 0 | 87.9 | 50.4 |
| 3 | 85.82 | 0 | 0 | 0 | 88.6 | 82.6 |
| 4 | 4.51 | 0 | 0 | 0 | 3.2 | 6.7 |
| 5 | 105.51 | 7.28 | 0 | 0 | 95.1 | 71.6 |
| 6 | 36.95 | 0 | 0 | 0 | 22.3 | 56.3 |
| 7 | 79.68 | 0.15 | 0 | 0 | 94.6 | 76.9 |
| 8 | 103.17 | 21.4 | 3.9 | 0 | 33.7 | 17.6 |
| Left frontal lobe lesions |  |  |  |  |  |  |
| 9 | 52.1 | 59 | 9.2 | 0.6 | 12.9 | 0 |
| 10 | 131.76 | 93.01 | 62.4 | 13.5 | 13.1 | 0.1 |
| 11 | 122.3 | 55.1 | 27.9 | 9.9 | 49.8 | 0 |
| 12 | 10.09 | 4.6 | 7 | 0 | 0 | 0 |
| 13 | 103.24 | 77.7 | 64.2 | 6.5 | 10.1 | 0 |

**Table 2.**
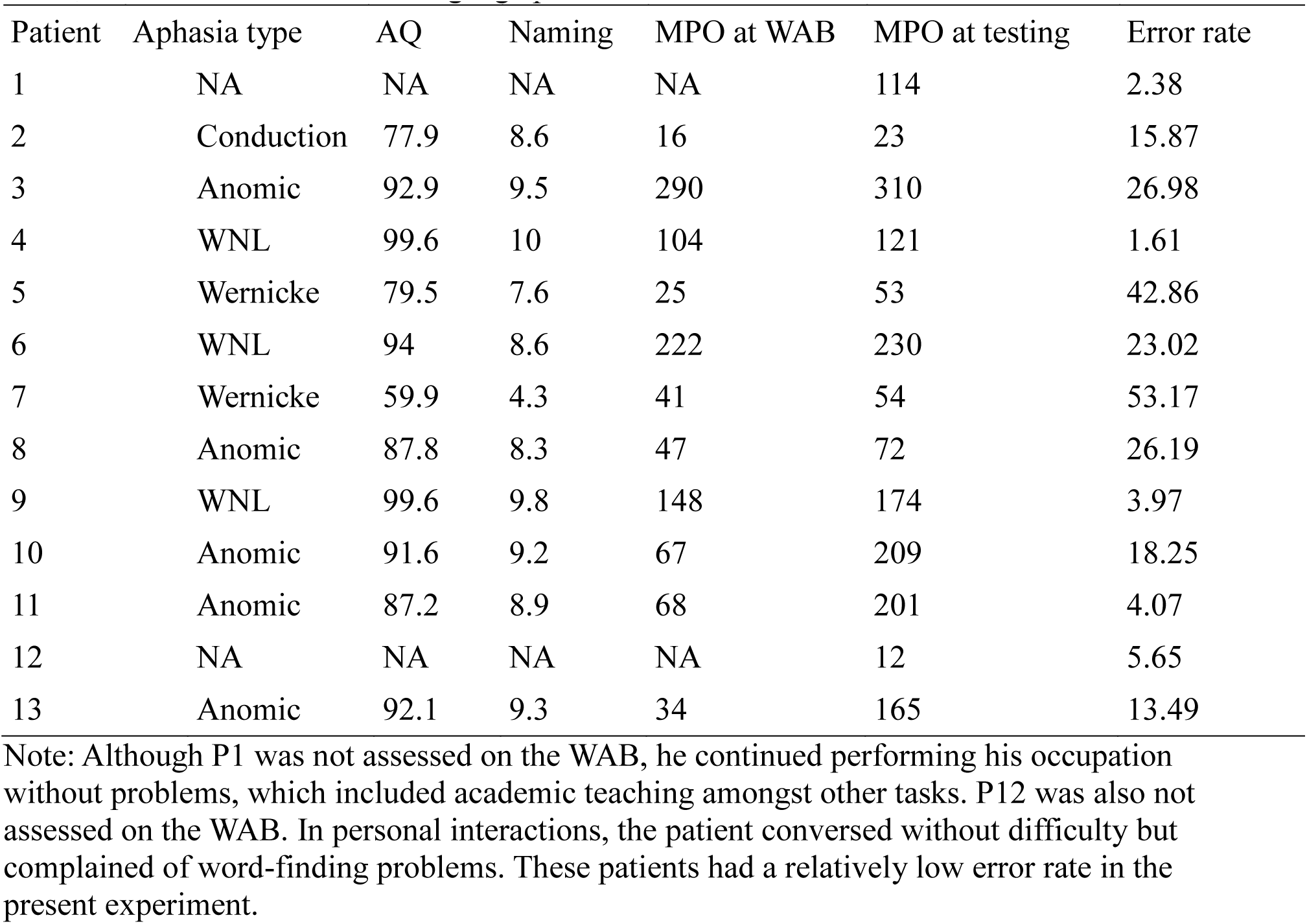
Language testing data from the Western Aphasia Battery (WAB). Naming = WAB Naming and Word Finding score (maximum = 10). Aphasia Quotient (AQ, maximum = 100). WNL = within normal limit. MPO = months post stroke onset. NA = not assessed on the WAB, see Methods section for language profile.

Individual lesions and overlap maps are shown for the frontal and temporal patients in Figure 2. In the frontal patients, the damage was centered mainly on the middle frontal gyrus and the most dorsal part of the inferior frontal gyrus (100% overlap). In the temporal patients, the damage was centered on the middle temporal gyrus (MTG, 100% overlap).

### Materials

Fifty-six color pictures were taken from the BOSS database (Brodeur, Dionne-Dostie, Montreuil, & Lepage, 2010). The pictures belonged to fourteen different semantic categories with four objects pertaining to each category (see Supplemental Materials). For each picture, distractor words were the picture name (“congruent” condition), from the same semantic category as the picture (“related”, distractor words were the names of the other category-coordinate pictures), or semantically and phonologically unrelated to the picture (“unrelated”, from recombining pictures with unrelated distractor words), as shown in Figure 1. Thus, all distractor words belonged to the response set. All participants saw each picture once in each condition. Pictures were presented on a white background on the center of the screen and distractors were presented in white, centered on the picture. The picture-word trials were randomized using Mix (van Casteren & Davis, 2006), with one unique list per participant. Participants were instructed to name the picture and to ignore the distractor word. Both speed and accuracy were emphasized.

**Figure 2.**
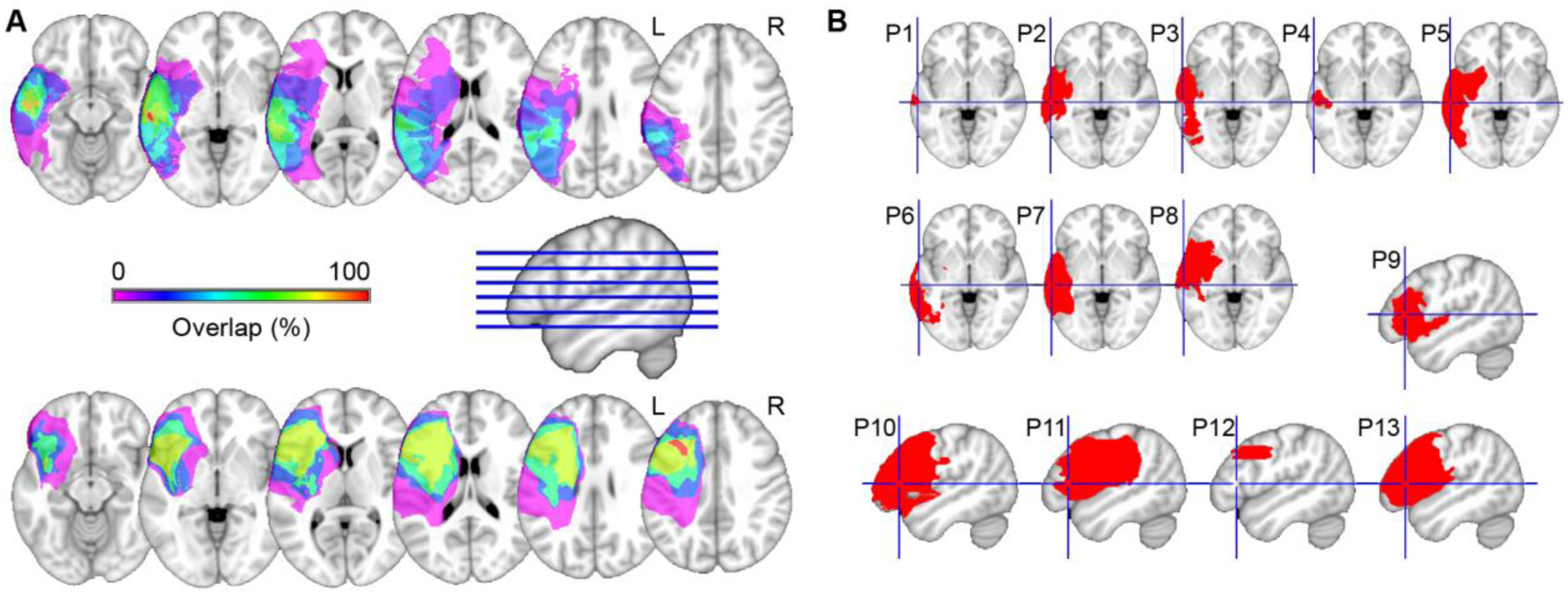
**A.** Lesion overlap map of the eight left temporal cortex patients (top) and of the five left prefrontal cortex patients (bottom). The color scale indicates the amount of overlap in lesion locations, with magenta indicating that only one patient had a lesion in that particular region (i.e., 0% overlap). **B.** Individual lesions on an axial slice (temporal patients, cross hairs indicate the middle temporal gyrus, MNI coordinates [-62, -25, -3]) and a sagittal slice (frontal patients, cross hairs indicate the left inferior frontal gyrus, MNI coordinates [-49, 26, 8]). L = left; R = right.

### Procedure

The presentation of stimuli and the recording of responses were controlled by Presentation Software (Neurobehavioral Systems). Participants were seated comfortably in front of a computer monitor. Vocal responses were recorded with a microphone. The experimenter evaluated the participants’ responses online. Trials began with a fixation cross presented for 1 s, followed by the presentation of the picture-word stimulus for 2 s. The intertrial interval varied between 1.25 and 2 s. No familiarization phase was used because we were concerned that patients would have different memory capacity that could confound the results.

### Analysis

Fourteen pictures were excluded from all analyses due to name agreement issues, yielding 42 trials for each participant per condition (see Supplemental Material for details). Responses containing dysfluencies or errors were coded in real time as incorrect and their corresponding trials excluded from all response time (RT) analyses. Naming RTs were calculated manually using Praat (Boersma & Weenink, 2013) before trials were separated by condition. The following responses were classified as errors: 1) the distractor word was named, 2) hesitations (e.g., the response started with filled pauses like “hum” or a poorly articulated initial phoneme), 3) no response was given, 4) phonological paraphasias, 5) a semantically related response (e.g., pictured bus, distractor “car”, response “truck”), 6) or another picture name was used than the expected name (e.g., “dish” for the picture of a bowl, “lime” for the picture of a lemon). This latter type of error was not considered a semantic error because it is possible that for the participant, that would be the correct label for the picture.

Single-trial RTs were analyzed with linear mixed-effects model and errors with mixed-effects logistic regression (Baayen, Davidson, & Bates, 2008), both with the same model structure. Models were fitted with the lme4-package (version 1.1.10; Bates, Maechler, Bolker, & Walker, 2015) in R (R Core Team, 2015). Single-trial RTs were log-transformed to reduce skewness and approach a normal distribution. In both models (referred to as “full model”), fixed effects for group (controls, temporal, and frontal patients) and distractor (related, unrelated, congruent) were included, as well as their interaction, and random intercepts for both participants and items. More complex models with random slope terms failed to converge. For “group”, controls were used as reference and for “distractor”, unrelated distractors were the reference. We additionally tested the semantic interference effect for each group separately with separate models (similar to the above, called “group model”). Due to the differences in language and lesion profile between frontal and temporal patients, the two groups are not compared to each other but rather to the reference control group. Significance of effects was obtained using the Satterthwaite approximation (lmerTest-package version 2.0.30, Kuznetsova, Brockhoff, & Christensen, 2016).

Hesitations were the most common error in all three groups (40% of the overall total and at least 33% of the total number of errors per group). Thus, we examined whether the number of hesitations was different for patient groups relative to controls with a Poisson regression model with fixed effect for group and random intercepts for participants. For each patient group, we also examined whether the error distribution differed between the related and unrelated conditions with Poisson regression models with fixed effect for distractor (related, unrelated) and random intercepts for participants.

## Results

Individual-averaged as well as group-averaged RTs and error rates are shown in Figure 3. Details on the statistics are shown in Tables 3 and 4.

**Figure 3.**
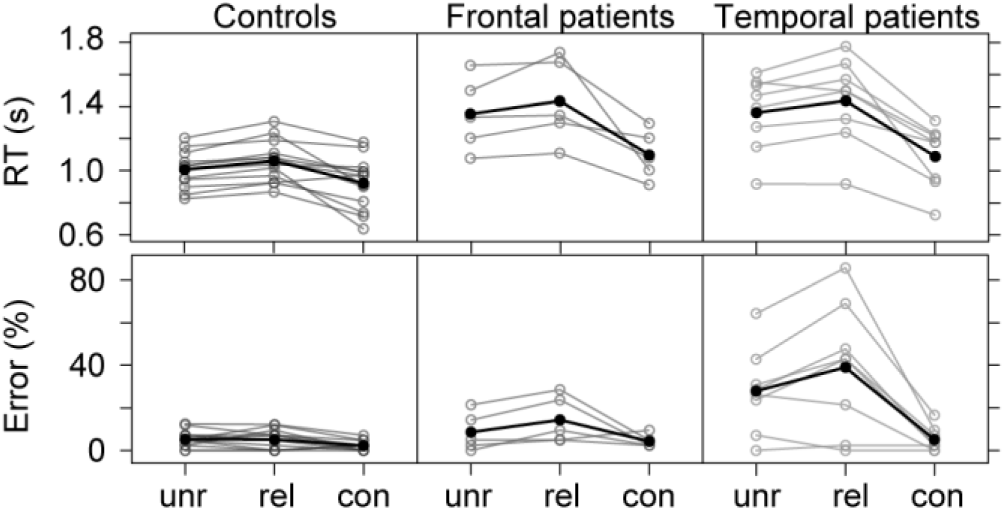
Individual-averaged (in brown, green, and purple) and group-averaged (in black) response times (RTs) and error rates for the three groups across conditions. Unr = unrelated; Rel = related; Con = congruent.

**Table 3.** Group-averaged response times in seconds and error rates (and standard deviations).

| Condition | Controls | Frontal | Temporal |
| --- | --- | --- | --- |
|  |  | Response times |  |
| Unrelated | 1.04 (0.22) | 1.45 (0.42) | 1.42 (0.38) |
| Related | 1.12 (0.26) | 1.49 (0.41) | 1.49 (0.37) |
| Congruent | 0.95 (0.24) | 1.19 (0.30) | 1.12 (0.28) |
|  |  | Error rates |  |
| Unrelated | 5.4 (3.9) | 8.6 (9.0) | 28.0 (20.0) |
| Related | 5.2 (4.5) | 14.4 (11.1) | 39.0 (30.1) |
| Congruent | 2.2 (2.3) | 4.3 (3.1) | 5.1 (5.8) |

**Table 4.** Results of the inferential statistics for the response times (RT, top) and error rates (bottom). Results obtained from the full model, unless stated otherwise. Results from the group models are indicated by an asterisk. SE = standard error

| RT effect | b | SE | t (df) | p |
| --- | --- | --- | --- | --- |
| Congruent vs unrelated | -.098 | .012 | -8.56 (2813) | < .001 |
| Related vs unrelated | .067 | .012 | 7.75 (2814) | < .001 |
| Frontal vs controls | .323 | .078 | 4.13 (24) | < .001 |
| Temporal vs controls | .308 | .067 | 4.60 (24) | < .001 |
| Related vs unrelated: controls* | .067 | .010 | 6.63 (974) | < .001 |
| Related vs unrelated: frontal* | .029 | .022 | 1.29 (323) | .198 |
| Related vs unrelated: temporal* | .039 | .018 | 2.22 (407) | .027 |
| Congruent vs unrelated: frontal vs controls | -.104 | .022 | -4.69 (2814) | < .001 |
| Related vs unrelated: frontal vs controls | -.036 | .022 | -1.60 (2817) | .109 |
| Congruent vs unrelated: temporal vs controls | -.151 | .020 | -7.68 (2816) | < .001 |
| Related vs unrelated: temporal vs controls | -.029 | .021 | -1.36 (2816) | .173 |
| Error rate effect | b | SE | z | p |
| Congruent vs unrelated | .968 | .341 | 2.822 | .005 |
| Related vs unrelated | .044 | .269 | .165 | .869 |
| Frontal vs controls | -.571 | .652 | -.876 | .381 |
| Temporal vs controls | -1.938 | .543 | -3.567 | < .001 |
| Related vs unrelated: controls* | .046 | .287 | .162 | .872 |
| Related vs unrelated: frontal* | -.682 | .327 | -2.087 | .037 |
| Related vs unrelated: temporal* | -.707 | .197 | -3.588 | < .001 |
| Congruent vs unrelated: frontal vs controls | -.163 | .543 | -.300 | .764 |
| Related vs unrelated: frontal vs controls | -.699 | .422 | -1.655 | .098 |
| Congruent vs unrelated: temporal vs controls | 1.465 | .455 | 3.216 | .001 |
| Related vs unrelated: temporal vs controls | -0.734 | .332 | -2.208 | .027 |

### RTs

Overall incongruency (unrelated vs congruent) and semantic interference (related vs unrelated) effects were observed (*ps* < .001). Patients were slower than controls (*ps* < .001). The incongruency effect was statistically larger for frontal and temporal patients than for controls (*ps* < .001). The semantic interference effect was significant for the controls (*p*< .001) and temporal patients (*p* = .027), but not for the frontal patients (*p* = .198). There was no evidence for a differential semantic interference effect between the controls and the two patient groups (*ps* > .109).

### Accuracy

Overall, the incongruency effect was significant (*p* = .005) but the semantic interference effect was not (*p* = .869). Temporal patients made more errors than controls and had a larger incongruency effect (*ps* =< .001). Temporal and frontal patients had a significant semantic interference effect (*ps* < .038). Temporal patients had a larger semantic interference effect than the controls (*p* = .027). No evidence was found for a difference between controls and frontal patients for the incongruency and semantic interference effects (*ps* > .098).

The error distribution is shown in Figure 4 and statistical results in Table 5. Temporal patients showed more hesitations than controls (99 vs 20, respectively, *p* < .001), whereas frontal patients and controls did not differ in the number of hesitations (28 vs 20, respectively). For the temporal patients, hesitations were more frequent with related than with unrelated distractors (58 vs 35, respectively, *p* = .017). The distributions were not significantly different for the frontal patients and controls (14 vs 10 and 12 vs 4, respectively).

**Figure 4.**
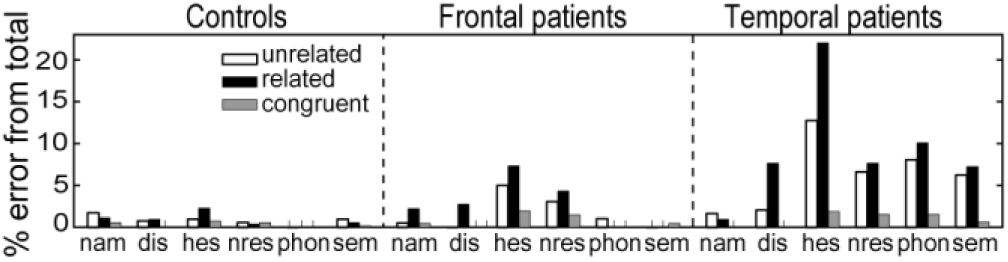
Error distribution in percentage from the total number of errors for the three groups across conditions. Nam = not the expected name; dis = distractor; hes = hesitation; nres = no response; phon = phonological paraphasia; sem = semantically related response. See Methods section for clarification.

**Table 5.** Results of the inferential statistics for the distribution of hesitations. SE = standard error

| Effect | b | SE | z value | p |
| --- | --- | --- | --- | --- |
| Frontal vs controls | 1.127 | .718 | 1.570 | .116 |
| Temporal vs controls | 1.185 | .604 | 3.62 | < .001 |
| Related vs unrelated: controls | 1.099 | .577 | 1.90 | .057 |
| Related vs unrelated: frontal | .337 | .414 | .813 | .416 |
| Related vs unrelated: temporal | .505 | .211 | 2.40 | .017 |

Aphasia quotient and lesion volume did not correlate with the magnitude of the semantic interference effect (for RTs and accuracy, *ps* > .504, see Supplemental Material).

## Discussion

Patients with left-lateral temporal and frontal lesions named pictures while ignoring semantically related, unrelated, and congruent visual distractors. The temporal patients had a significant semantic interference effect both in the error rates and in the RTs. They also had an increased semantic interference effect in the error rates relative to controls. Hesitations in language production have been related to difficulties in lexical selection (Goldman-Eisler, 1968). The analysis of hesitations corroborated the findings of the semantic interference effect in the temporal patients in that more hesitations were present in the responses for the related than for the unrelated condition.

In the RTs, we observed an increased incongruency effect in the patients relative to controls. For the temporal patients, the increased incongruency effect was additionally found in the error rates. The congruent condition was included in the present study to maximize the interference from distractors (cf. Lowe & Mitterer, 1982). Given the theoretical relevance of the *semantic interference effect*, we focus the remainder of the discussion on this effect.

Theories of lexical selection differ in the extent to which frontal-cortex structures, or components external to the lexical system, are involved in the selection process. Some models argue that selection depends on lateral PFC structures and that representations compete in the ventrolateral PFC (e.g., Snyder et al., 2010; Snyder, Banich, & Munakata, 2014). In WEAVER++, the activation of target nodes in the network are enhanced selectively until goals in working memory are achieved (Roelofs, 1992, 2003). These enhancements have been linked to medial frontal structures, such as the anterior cingulate cortex (Roelofs & Hagoort, 2002), but lexical representations “compete” in the left temporal lobe (Roelofs, 2014). In the two-step model of lexical access (Dell, Schwartz, Martin, Saffran, & Gagnon, 1997), a jolt of activation to semantic features gives rise to spreading activation through the network. Lexical selection is concluded with the node with the highest activation being selected. In this case, selection is an intrinsic part of the lexical-access process.

We have not obtained evidence that a top-down control process exercised by the ventrolateral PFC is necessary for resolving competition from semantically related distractor words. Otherwise, we should have observed a disproportional semantic interference effect for the PFC patients relative to controls (see for similar findings using other paradigms Britt, Ferrara, & Mirman, 2016; Riès, Karzmark, Navarrete, Knight, & Dronkers, 2015). However, this conclusion is somewhat limited by the fact that in the present study, as well as in Piai et al. (2016), no semantic interference effect was observed on the group level for patients with left PFC damage. Top-down regulation of lexical selection in the presence of competing semantic distractors might be subserved by a different region, such as superior medial-frontal structures (Roelofs & Hagoort, 2002). This account finds support in previous neuroimaging studies of picture-word interference (Piai et al., 2013, 2014), and neuroimaging and neuropsychological findings on verbal tasks involving control (Alario, Chainay, Lehericy, & Cohen, 2006; Derrfuss, Brass, Neumann, & von Cramon, 2005; Stuss, Floden, Alexander, Levine, & Katz, 2001). However, lesion-symptom studies of picture-word interference involving medial frontal structures are lacking. Future studies are needed to clarify the role of frontal-cortex structures in lexical selection in the presence of competing semantic distractors.

Regarding the left temporal cortex, this is the first study to examine the semantic interference effect in a group of patients with well-characterized lesions. We found that patients with left temporal lesions (overlapping fully in the mid portion of the MTG) made more errors with related than with unrelated distractors. Various types of errors in word production are thought to emerge from the incorrect selection of words (Schwartz, Dell, Martin, Gahl, & Sobel, 2006; Roelofs, 1992). A large literature suggests a critical role for the MTG in naming (Baldo, Arévalo, Patterson, & Dronkers, 2013; Schwartz et al., 2009) and the mid portion of the MTG in particular is thought to subserve word activation and selection (Indefrey & Levelt, 2004). Lexical selection takes place once the activation of the target node exceeds that of all other competitors (by some critical amount, e.g., Roelofs, 1992). Left MTG lesions likely introduce noise to the activation of representations of both target and competitors, making these representations more similar. Accordingly, selection difficulty, as in the case of hesitations, and selection errors are more likely to occur with noisy competing representations that do not show sufficient activation differences. Our results are more consistent with models in which lexical selection is an intrinsic part of the lexical activation dynamics, tightly related to the left temporal lobe.

In conclusion, the left temporal lobe is a necessary structure for lexical selection in word production. Following the view that conceptually driven word retrieval involves activation of candidate words and competitive selection of the intended word from this set, we argue that left middle temporal lesions affect the lexical activation component. A deficit in this component makes lexical selection more susceptible to errors.

## Supplemental material

### Materials

The pictures used in the experiment, as well as name agreement norms (i.e., proportion of participants who gave the target name relative to the number of participants who named the picture) from the BOSS database (Brodeur et al., 2010) are shown in Table S1.

**Table S1.** Pictures used in the experiment and their name agreement from the BOSS database and from the present experiment.

| Picture | Target name in exp | Name agreement from BOSS (%) | Used for analyses | Name agreement from controls' first responses (%) | Remark |
| --- | --- | --- | --- | --- | --- |
| pants | pants | 79 | no | 100 | excluded because "shorts" was consistently named "pants" |
| scarf | scarf | 95 | yes | 100 |  |
| shirt01 | shirt | 44 | no | 61.5 | other names: jacket, coat |
| shorts01 | shorts | 74 | no | 30.8 | consistently named "pants" |
| drill01b | drill | 47 | yes | 100 |  |
| hammer01 | hammer | 100 | yes | 100 |  |
| pliers02b | pliers | 85 | yes | 100 |  |
| saw02b | saw | 88 | yes | 100 |  |
| cow | cow | 93 | yes | 100 |  |
| horse | horse | 100 | yes | 100 |  |
| pig | pig | 90 | yes | 100 |  |
| sheep | sheep | 93 | yes | 100 |  |
| cannon | cannon | 93 | yes | 100 |  |
| revolver | gun | 43 | yes | 100 |  |
| spear02 | spear | 75 | no | 23.1 | other names: arrow, paintbrush |
| sword01 | sword | 81 | yes | 100 |  |
| bowl01 | bowl | 67 | yes | 100 |  |
| jug | jug | 31 | no | 23.1 | consistently named "pitcher" |
| mug05 | mug | 50 | no | 30.8 | consistently named "cup" |
| plate01b | plate | 92 | yes | 92.3 |  |
| arm | arm | 71 | yes | 100 |  |
| ear | ear | 83 | yes | 100 |  |
| leg | leg | 60 | yes | 92.3 |  |
| nose | nose | 88 | yes | 100 |  |
| bed | bed | 81 | yes | 100 |  |
| chair | chair | 88 | yes | 100 |  |
| dresser02 | dresser | 49 | no | 61.5 | other names: bureau,<br>chest, table |
| table01 | table | 81 | yes | 100 |  |
| apple07 | apple | 95 | yes | 100 |  |
| banana01 | banana | 100 | yes | 100 |  |
| lemon02 | lemon | 100 | yes | 100 |  |
| pear01 | pear | 100 | yes | 100 |  |
| acousticguitar02 | guitar | 71 | yes | 100 |  |
| drum01 | drum | 51 | yes | 92.3 |  |
| grandpiano | piano | 55 | yes | 100 |  |
| violin | violin | 97 | yes | 100 |  |
| barn | barn | 71 | no | 61.5 | other names: shed,<br>house |
| castle | castle | 61 | yes | 69.2 |  |
| church | church | 81 | yes | 92.3 |  |
| shed02 | shed | 60 | no | 38.5 | other name: garage |
| binder03b | binder | 68 | no | 30.8 | other names: book,<br>notebook |
| book01b | book | 90 | yes | 100 |  |
| pen04b | pen | 90 | yes | 100 |  |
| pencil01 | pencil | 97 | yes | 100 |  |
| airplane* | airplane | NA | yes | 100 |  |
| bicycle | bicycle | 43 | yes | 100 |  |
| bus | bus | 57 | yes | 100 |  |
| car | car | 81 | yes | 100 |  |
| bracelet01 | bracelet | 62 | no | 69.2 | other name: necklace,<br>excluded because<br>"necklace" was<br>another picture in the<br>experiment |
| necklace | necklace | 81 | no | 61.5 | other name: bracelet |
| ring01 | ring | 100 | no | 61.5 | other name: bracelet |
| watch02a | watch | 87 | no | 61.5 | other name: bracelet |
| broccoli01a | broccoli | 97 | yes | 100 |  |
| cucumber | cucumber | 95 | yes | 100 |  |
| lettuce | lettuce | 68 | yes | 92.3 |  |
| pumpkin | Pumpkin | 98 | yes | 100 |  |

\* The picture “airplane” was taken from our own database. Exp = experiment

We calculated the proportion of control participants who named a given picture with its target name at its first presentation, that is, the first time participants encounter each picture in the experiment. The distribution of name-agreement values is shown in Figure S1 for the pictures included in the analyses and the pictures not included in the analyses. The actual name-agreement values are shown in Table S1 above. As can be seen, all but one item kept for analysis had name-agreement values above 90%. Two pictures had name agreement above 70% (“pants” and “bracelet”), but they were not kept in the analyses because other two pictures (“shorts” and “necklace”) were often named “pants” and “bracelet”.

**Figure S1.**
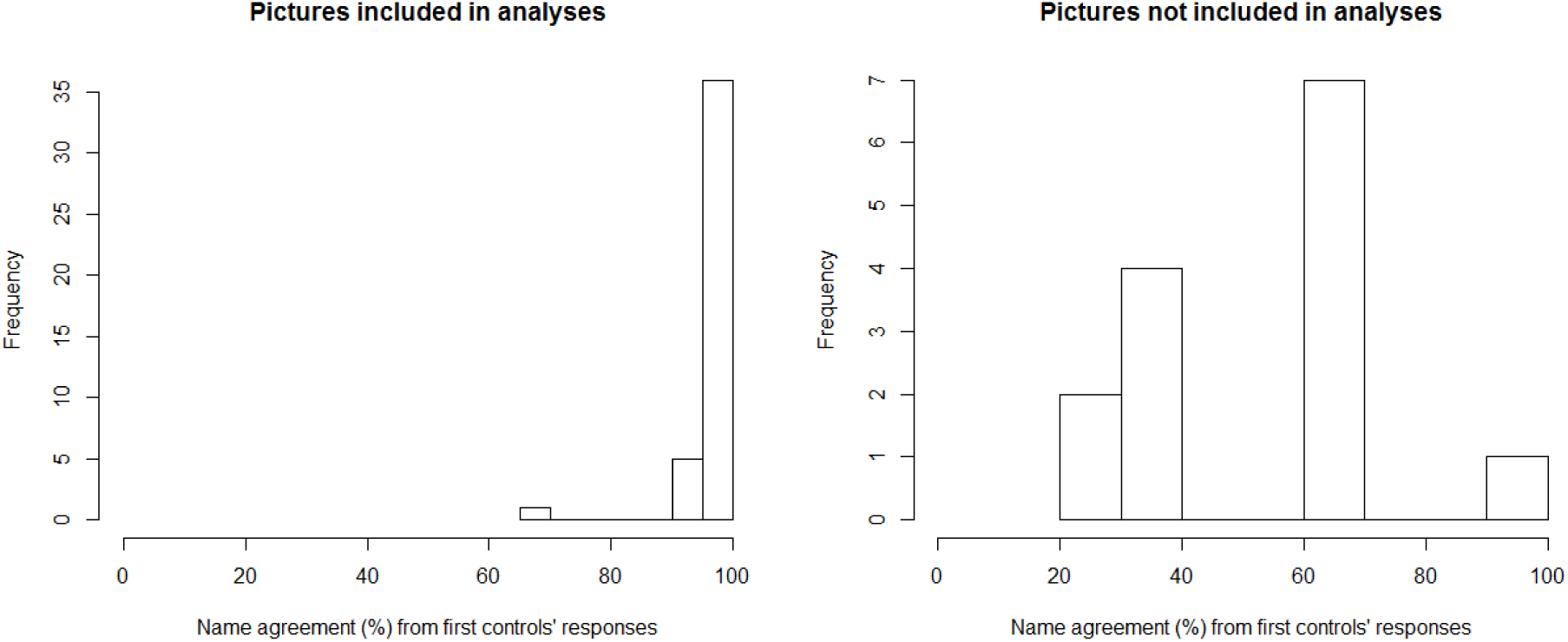
Distribution of name-agreement values, calculated from the first presentation of each item in the experiment for the control participants, for the picture items included in the analyses (left) and for those excluded from analyses (right).

We also ran the same analyses on the RTs as reported in the main text after excluding trials in an item-wise fashion. That is, if a participant named, for example, the picture item “lemon” incorrectly once, all other trials corresponding to the picture item “lemon” were also excluded for this participant, even if the other responses were correct.

Overall incongruency (unrelated vs congruent) and semantic interference (related vs unrelated) effects were observed (*ps* < .001). Patients were slower than controls (*ps* < .001). The incongruency effect was statistically larger for frontal and temporal patients than for controls (*ps* < .001). The semantic interference effect was significant for the controls (*p*< .001) and temporal patients (*p* =.037), but not for the frontal patients (*p* =.124). There was no evidence for a differential semantic interference effect between the controls and the two patient groups (*ps* > .134). Details on the statistics are shown in Table S2. Thus, the pattern of results was the same as reported in the main text.

**Table S2.**
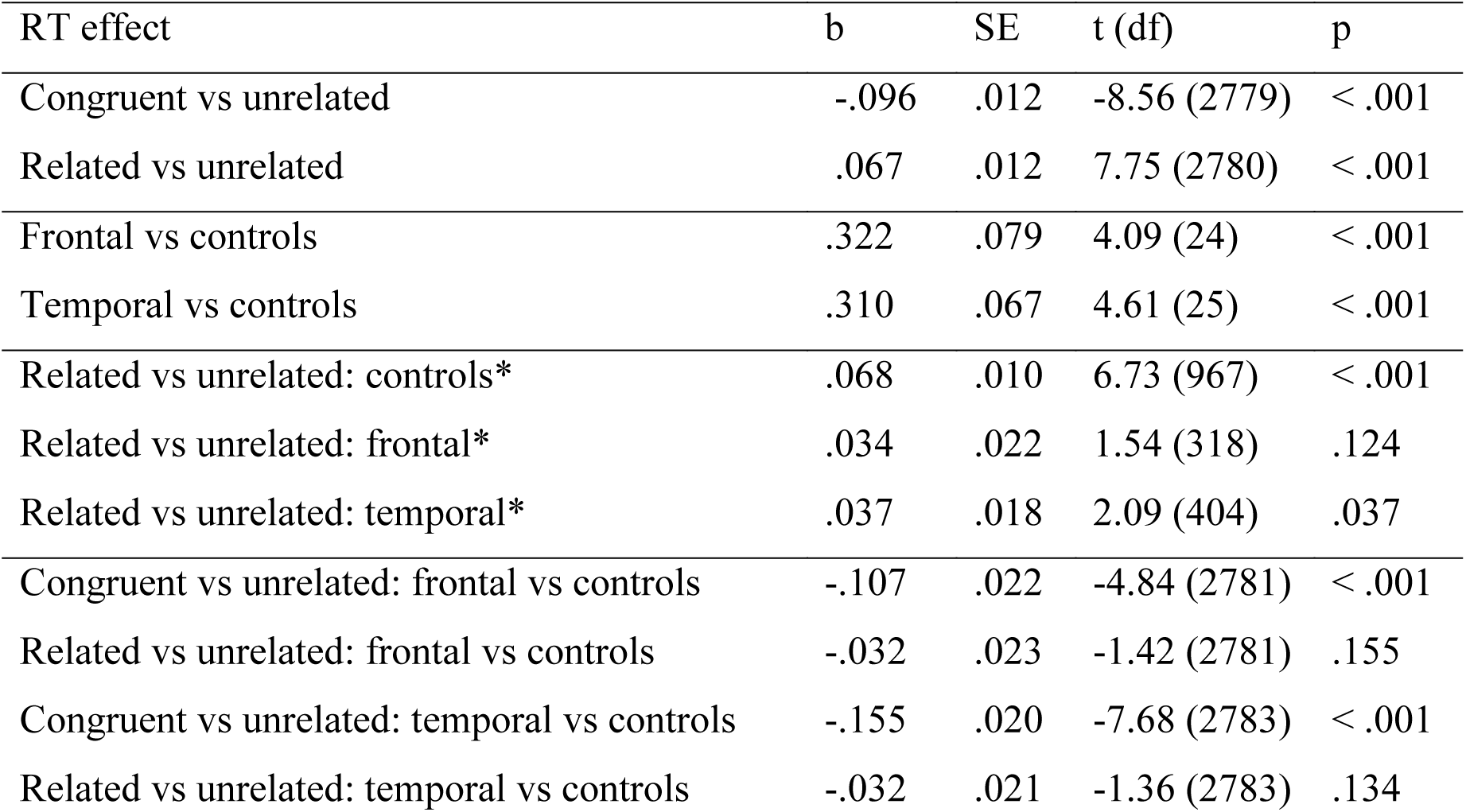
Results of the inferential statistics for the response times (RT, top) and error rates (bottom). Results obtained from the full model, unless stated otherwise. Results from the group models are indicated by an asterisk. SE = standard error.

| RT effect | b | SE | t (df) | p |
| --- | --- | --- | --- | --- |
| Congruent vs unrelated | -.096 | .012 | -8.56 (2779) | < .001 |
| Related vs unrelated | .067 | .012 | 7.75 (2780) | < .001 |
| Frontal vs controls | .322 | .079 | 4.09 (24) | < .001 |
| Temporal vs controls | .310 | .067 | 4.61 (25) | < .001 |
| Related vs unrelated: controls* | .068 | .010 | 6.73 (967) | < .001 |
| Related vs unrelated: frontal* | .034 | .022 | 1.54 (318) | .124 |
| Related vs unrelated: temporal* | .037 | .018 | 2.09 (404) | .037 |
| Congruent vs unrelated: frontal vs controls | -.107 | .022 | -4.84 (2781) | < .001 |
| Related vs unrelated: frontal vs controls | -.032 | .023 | -1.42 (2781) | .155 |
| Congruent vs unrelated: temporal vs controls | -.155 | .020 | -7.68 (2783) | < .001 |
| Related vs unrelated: temporal vs controls | -.032 | .021 | -1.36 (2783) | .134 |

Figure S2 shows scatterplots of the relationships between aphasia quotient or lesion volume and the semantic interference effect in response times (RTs) and error rates.

**Figure S2.**
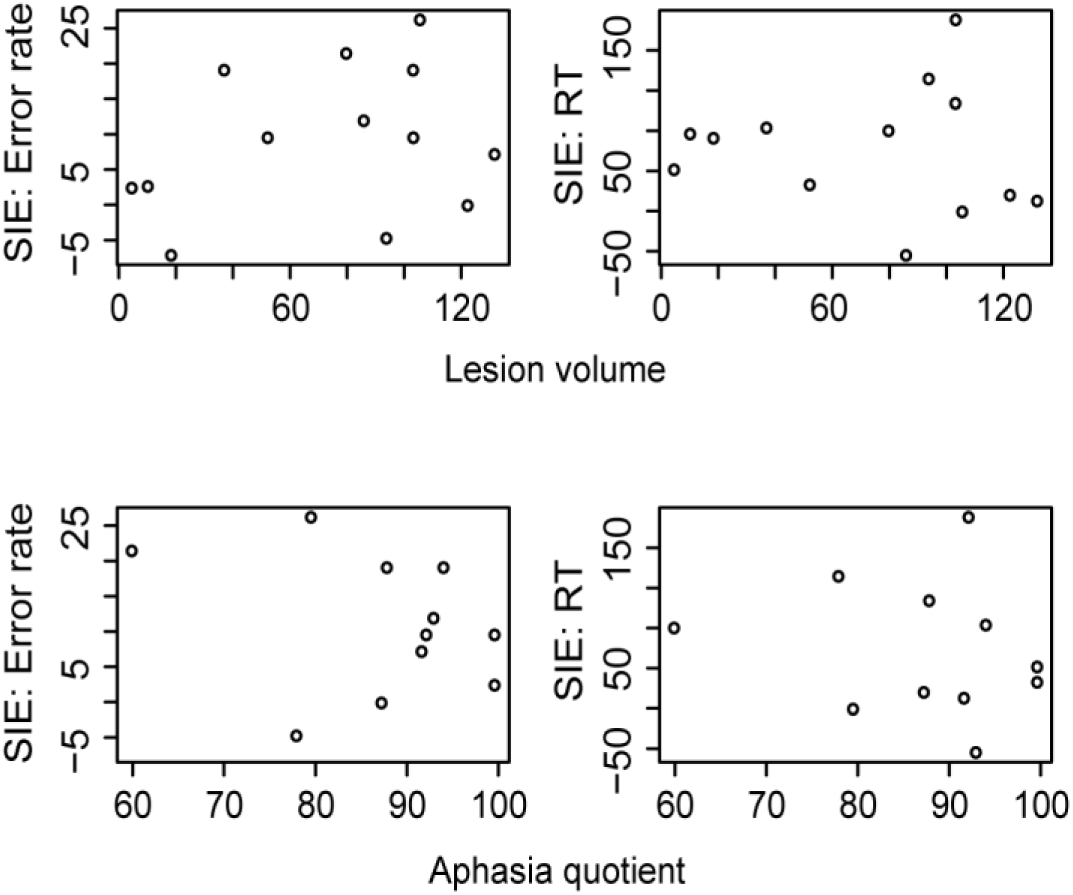
Scatterplots for lesion volume (top) and aphasia quotient (bottom) and the semantic interference effect (SIE) in error rates (left) and response times (RT, right). Lesion volume did not correlate with the magnitude of the semantic interference effect in the RTs (Spearman’s *rho* = -.187, *S* = 432, *p* =.541) nor in the error rates (Spearman’s *rho* =.204, *S* = 290, *p* =.504). Aphasia quotient (available for 11 patients) did not correlate with the magnitude of the semantic interference effect in the RTs (Spearman’s *rho* = -.105, *S* = 243, *p* =.759) nor in the error rates (Spearman’s *rho* = -.149, *S* = 253, *p* =.663).

